# Preserving spiking and EEG in detailed models of human cortical microcircuits with morphological reductions

**DOI:** 10.64898/2026.09.18.752744

**Authors:** Alexandre Guet-McCreight, Etay Hay

## Abstract

Biophysically detailed models of human cortical microcircuit provide key insights into brain function and disease biomarkers, yet their computational cost lengthens simulation runtime and limits scalability. To address this, we systematically evaluated recent simplification methods for morphological reduction and synapse merging in terms of preserving spiking and electroencephalographic (EEG) properties of human cortical microcircuit models. At the single-cell level, reduced compartmental models accurately reproduced spiking behaviors. However, at the microcircuit level, while achieving a 5-fold speed-up, morphological reduction failed to preserve spectral dynamics of spiking and EEG despite maintaining average baseline and response firing rates. Conversely, applying synapse merging alone on the full morphology successfully reproduced both spiking activity and EEG dynamics, when constrained to ≤ 50 µm spatial intervals, and yielded a 2-fold speed-up. These results were consistent across both cortical layer 2/3 and layer 5 models, and in depression models with reduced somatostatin interneuron inhibition. Our study identifies model simplification methods that speed up cortical microcircuit simulations while preserving the accuracy of the simulated brain signals.

## Introduction

Detailed microcircuit models provide insights into how cellular and circuit mechanisms shape cortical computation and non-invasive brain signals, like electroencephalographic (EEG) recordings^1–4^. Despite their utility, the computational cost of simulating detailed morphologies lengthens runtimes, which limits their application in large-scale network simulations^5,6^ and clinical-scale modeling^7^. Recent model reduction methods collapsed dendritic trees into simplified structures and merged synapses onto the resulting compartments^8–10^. While the studies showed that the reduction maintains some properties of the single neurons, the impact on circuit activity and spatial current dipoles responsible for EEG dynamics^11–13^ has not been assessed.

Model reduction methodologies aim to simplify high-dimensional neuron models while preserving biophysical fidelity. Earlier studies used analytical geometric reduction to collapse dendritic trees by conserving surface area and axial resistance^9^ and employing topological template-based simplifications^10^. While other analytical frameworks, such as linear least-squares solutions^14^, also reduce complex morphologies to a minimal number of compartments while retaining the model active non-linear dendritic properties, they did not address preserving the input-output properties through synaptic integration. A recent analytical method maps complex dendrites and synapses to merge onto simplified multi-cylindrical structures via transfer impedance conservation^8^. This method preserved the input and output properties of complex neuron morphological models with realistic numbers of synapses^8^, although it has not been tested on preserving activity and signals from microcircuits.

Morphological models of neural circuits have been used extensively to bridge the gap between microscopic cellular activity and macroscopic non-invasive brain signals, such as LFP, EEG and MEG, by simulating the combined electrical currents generated across neuronal compartments during cortical activity^13,15–19^. We previously developed biophysically detailed models of human cortical microcircuits^1^, which integrated cellular and circuit human data, and used these to identify EEG-based cellular biomarkers of depression^2^, aging^4^, and drug efficacy^3,7,20^. The models included different layers of human cortical microcircuits with distinct activity and oscillatory dynamics^21^. In this work we use the human cortical microcircuit models to assess the recent morphological reduction and synapse merging methods in terms of preserving microcircuit spiking activity and EEG, across different neuron types, cortical layers and disease models of depression.

## Results

To reduce our layer 2/3 microcircuit models, we started by adopting a morphological reduction method that preserves the transfer impedance from each dendritic compartment to the soma (**Fig. 1A**). We further accounted for dendritic mechanisms by integrating this approach with a method for scaling the ion channel conductance values and membrane capacitances by the resulting changes in dendritic surface areas (**Fig. 1A**). The reduced morphological models sufficiently preserved somatic spiking features in response to step current, spike frequency-input curve, and sag amplitudes, for each cell type in the microcircuit (**Fig. 1B-E**), with small relative differences between the full and reduced models (< 8% for most features, except for -11% Pyr spike after-hyperpolarization depth and -10% PV spike rate).

**Figure 1.**
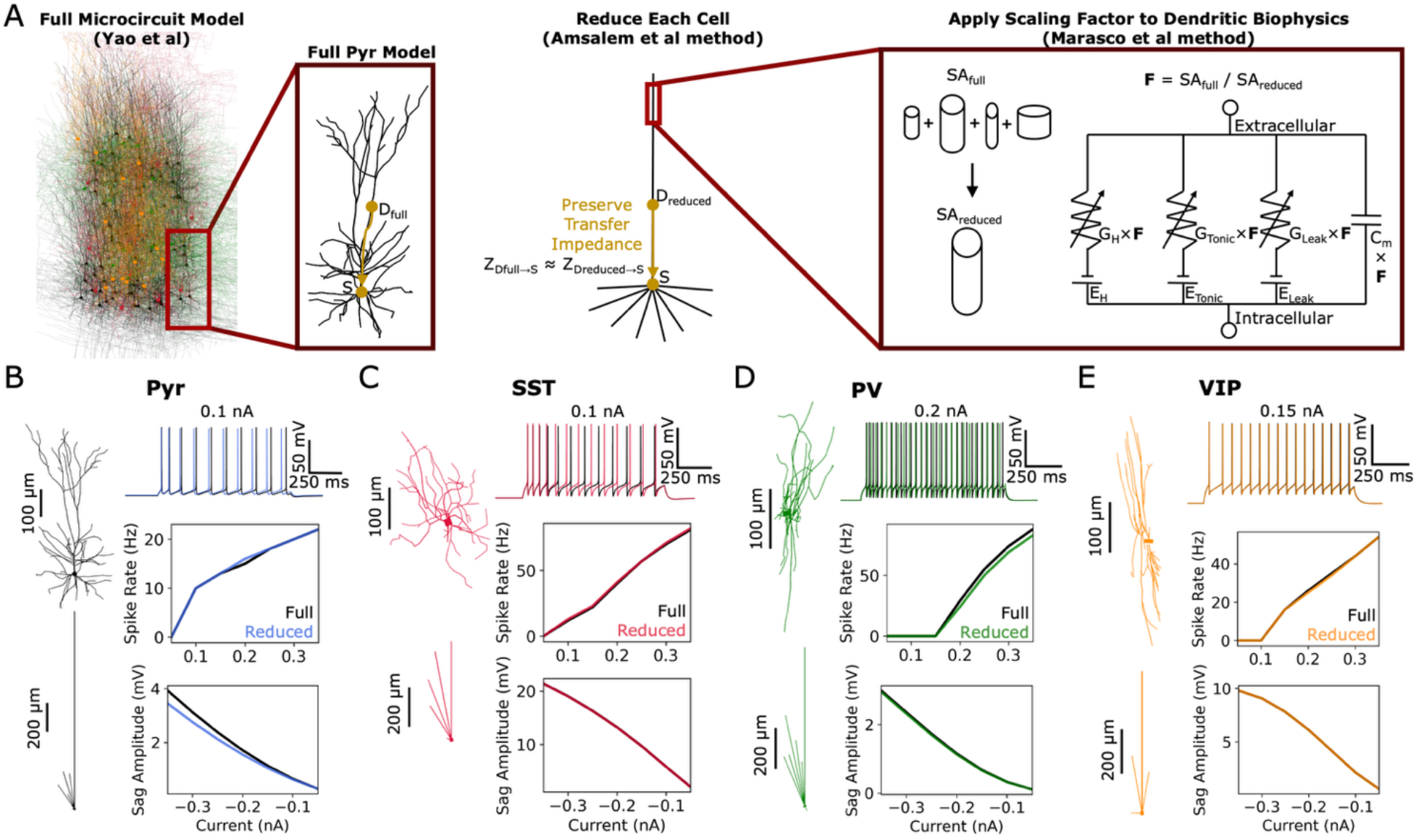
Reduced human L2/3 neuron models replicate full model excitability. A. Illustration of morphological and synaptic reduction method. Beginning with the full layer 2/3 human microcircuit model (Yao et al 2022), step 1: reduce the morphological complexity and synaptic number in each cell in the microcircuit, preserving transfer impedance (*Z*) from dendrites (*D*) to soma (*S*), and remapping synaptic point processes to the corresponding reduced dendritic compartments (Amsalem et al method); Step 2: scale the dendritic conductance (*G*) and membrane capacitance (*Cm*) in the reduced dendritic compartments according to the ratio (*F*) of surface area (*SA*) change from full to reduced morphology (Marasco et al method). B-E. Left: full and reduced morphologies for each neuron type (B: pyramidal – Pyr, C: somatostatin – SST, D: parvalbumin – E: PV, vasoactive intestinal polypeptide – VIP). Right: comparison of example step-current 4 spiking response (top), spike frequency-input curve (middle), and sag amplitude (bottom) for each neuron type with full (black) or reduced (color) morphology.

We next assessed implementing the reduced morphology models into microcircuits, where synapses get merged onto the reduced compartments (**Fig. 2A**). For this, we compared the full and reduced microcircuit models at baseline and stimulus response (**Fig. 2B**) and found Pyr neuron spike rates were unchanged (baseline firing rate, full: 0.76 ± 0.04 Hz, reduced: 0.72 ± 0.04 Hz; response firing rates, full: 2.53 ± 0.76 Hz, reduced: 2.36 ± 0.37 Hz; **Fig. 2C**), as well as interneuron spike rates (SST: 5.72 ± 0.21 Hz vs. 5.66 ± 0.23 Hz, PV: 10.07 ± 0.43 Hz vs. 10.10 ± 0.46 Hz, VIP: 3.50 ± 0.27 Hz vs. 3.47 ± 0.28 Hz; **Fig. 2D**).

**Figure 2.**
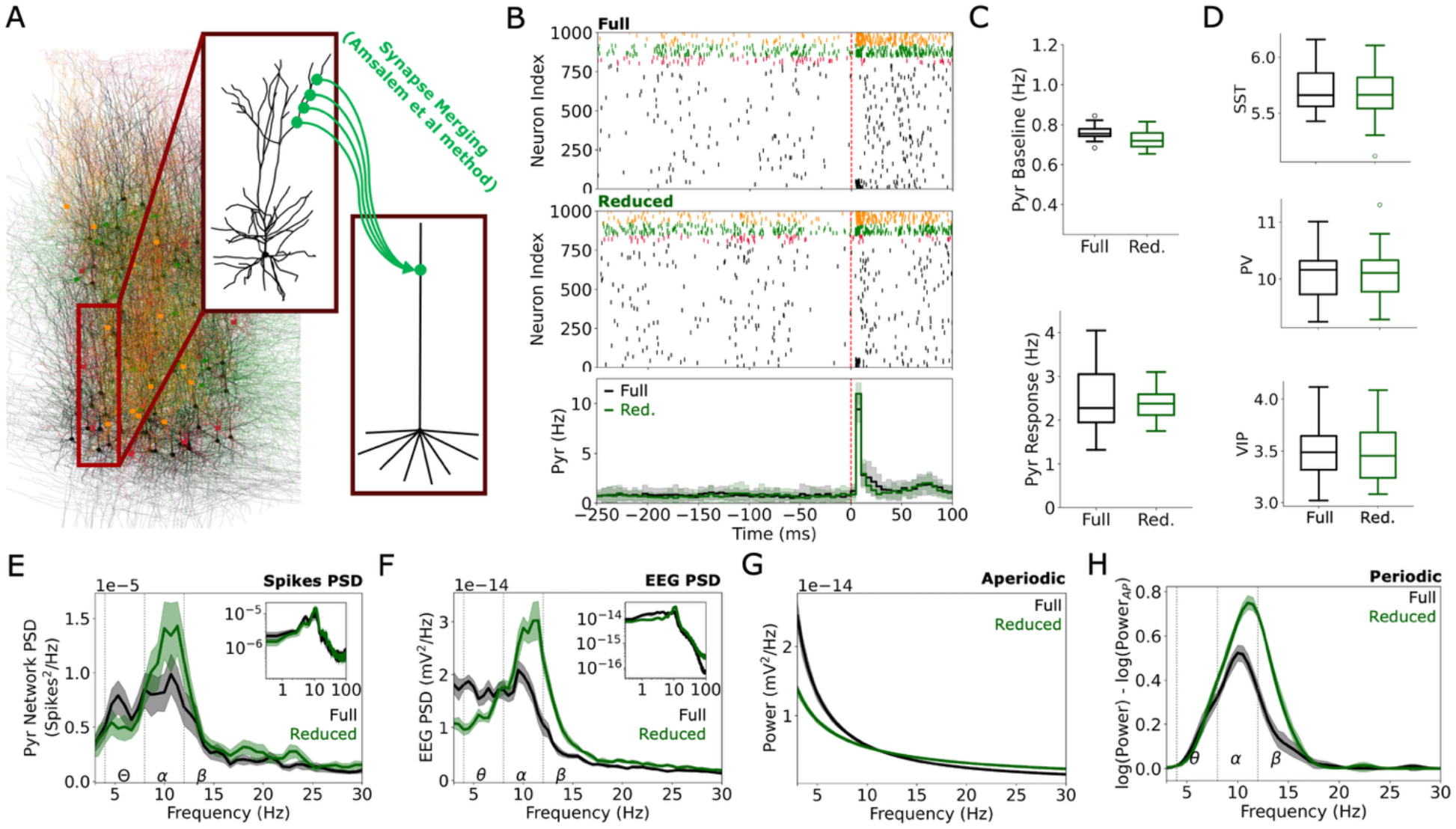
Reduced human L2/3 cortical microcircuits preserve spiking of the full models but not spectral features and EEG. **A**. Schematic of synapse merging following microcircuit reduction using the method of Amsalem et al. **B**. Example raster plots for full (top) and reduced (middle) L2/3 microcircuits. Bottom: peri-stimulus time histograms of spiking for Pyr neurons in full (black) and reduced (green) microcircuits. Standard deviations are shown in shaded areas. **C**. Baseline and response spike rates of Pyr neurons in the full and reduced microcircuits (n = 20 microcircuits each). **D**. Baseline spike rates of the different interneuron populations in the full and reduced microcircuits. **E**. Mean spikes PSD of Pyr neurons for full and reduced microcircuits. Inset: PSD in log scale, illustrating the 1/f relationship between power and log frequency. **F**. PSD of simulated EEG of full and reduced microcircuits (bootstrapped mean, and 95% confidence intervals). Inset: PSD in log scale. **G-H**. Aperiodic (G) and periodic (H) components of the PSD (n = 10 microcircuits each).

We further analyzed the spiking power spectral dynamics of Pyr neuron population activity and primarily found a 43.4% increase in α power for the reduced model (full: 3.36 × 10^5^ ± 0.84 × 10^5^ spikes^2^; reduced: 4.82 × 10^5^ ± 1.71 × 10^5^ spikes^2^; *p* < 0.05, Cohen’s *d* = 1.0; **Fig. 2E**). Mismatches were more pronounced and broader in the simulated EEG power spectra, with a 25.6% decrease in θ power (0.0679 ± 0.0044 nV^2^ vs. 0.0505 ± 0.0036 nV^2^, *p* < 0.05, Cohen’s *d* = -4.0, **Fig. 2F**), a 49.1% increase in α power (0.0672 ± 0.0060 nV^2^ vs. 0.1002 ± 0.0093 nV^2^, *p* < 0.05, Cohen’s *d* = 4.0), and 69.2% increase in β power (0.0383 ± 0.0022 nV^2^ vs. 0.0648 ± 0.0043 nV^2^, *p* < 0.05, Cohen’s *d* = 7.3). Mismatches in the EEG power spectra could also be localized to deviations in both the aperiodic component (offset: -13.04 ± 0.10 a.u. vs. -13.5 ± 0.06 a.u., *p* < 0.05, Cohen’s *d* = -5.2; exponent: 1.20 ± 0.08 vs. 0.77 ± 0.04, *p* < 0.05, Cohen’s *d* = -6.4, **Fig. 2G**) and periodic component (α peak: 0.51 ± 0.10 a.u. vs. 0.73 ± 0.10 a.u., *p* < 0.05, Cohen’s *d* = 2.1; α width: 2.78 ± 0.89 Hz vs. 4.04 ± 1.12 Hz, *p* < 0.05, Cohen’s *d* = 1.2, **Fig. 2H**).

We next examined an alternative reduction method that may better preserve both spiking and EEG dynamics, by adopting the synapse merging method above without the morphological reduction step, at different spatial intervals of merging on the full morphology models (**Fig. 3A**). We note that merging the synapse point processes without moving them spatially (M0), while still a computational load optimization, is mathematically equivalent to our full microcircuit models, and thus the comparisons we show below are relative to this case. Merging synapses onto their nearest 50 μm dendritic interval from the soma (M50) preserved both baseline and response Pyr spike rates (baseline: 0.77 ± 0.04 Hz; response: 2.29 ± 0.49 Hz; **Fig. 3B**). Increasing the spatial intervals further to 100 μm (M100) introduced differences in the pre- and post-stimulus Pyr spike rates (pre: 0.86 ± 0.05 Hz, *p* < 0.05, Cohen’s *d* = 2.0; post: 5.02 ± 1.19 Hz, *p* < 0.05, Cohen’s *d* = 1.8; **Fig. 3B**). The interneuron pre-stimulus spike rates also remained unchanged for the M50 condition (SST: 5.81 ± 0.17 Hz, PV: 10.16 ± 0.42 Hz, VIP: 3.55 ± 0.32 Hz) and differed in the M100 condition (SST: 6.20 ± 0.22 Hz, *p* < 0.05, Cohen’s *d* = 2.1, PV: 10.77 ± 0.48 Hz, *p* < 0.05, Cohen’s *d* = 1.3, VIP: 3.99 ± 0.28 Hz, *p* < 0.05, Cohen’s *d* = 1.5; **Fig. 3C**).

**Figure 3.**
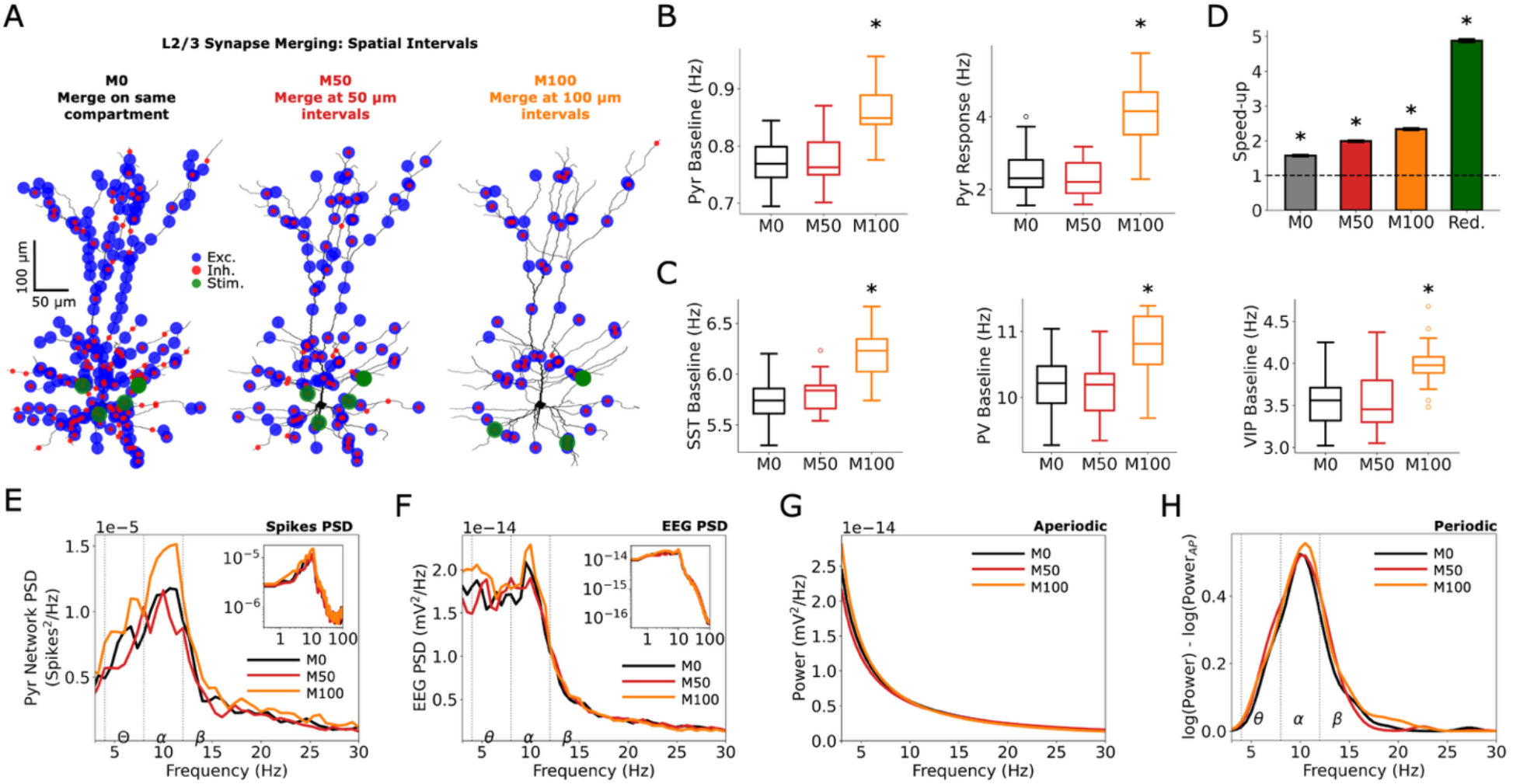
Spatially merging synapses speeds-up simulation runtime while preserving spiking and EEG. **A**. Example merging of L2/3 Pyr neuron synapse locations with different merging spatial resolutions. **B**. Baseline (left) and response (right) spike rates of Pyr neuron for each merging spatial resolution. **C**. Baseline spike rates of the different interneuron types for each merging spatial resolution **D**. Mean speed-ups relative to the full microcircuit model (4.5s-long simulation, without EEG recordings), for the full microcircuit with synapses merged across different spatial resolutions, and the reduced morphology microcircuit from fig 1 (green) (n = 20 microcircuits each). **E**. Mean spikes PSD of Pyr neurons for each spatial resolution. Inset: PSD in log scale. **F**. Mean PSD of simulated EEG for each spatial resolution. Inset – PSD in log scale. **G-H**. Aperiodic (G) and periodic (H) components of the PSD (n = 10 microcircuits each). Asterisks denote significant differences (*p* < 0.05; Cohen’s *d* > 1) compared to the full microcircuit.

The spiking power spectra of the M50 model did not exhibit differences from that of M0, but for the M100 models there was increased α power (5.45 × 10^5^ ± 1.90 × 10^5^ spikes^2^, *p* < 0.05, Cohen’s *d* = 0.66; **Fig. 3E**). In the EEG power spectra, only the M100 exhibited an increase in θ power (0.0752 ± 0.0069 nV^2^, *p* < 0.05, Cohen’s *d* = 1.2; **Fig. 3F**). Although mismatches in the aperiodic and periodic components were visually marginal, we do report mismatches for both the M50 and M100 models (M50 α width: 4.06 ± 1.01 Hz, *p* < 0.05, Cohen’s *d* = 1.3; M100 exponent: 1.31 ± 0.11, *p* < 0.05, Cohen’s *d* = 1.1; **Fig. 3G-H**). We also assessed the speed-up of morphological reductions and synapse merging at different spatial intervals, and all cases generated significant speed-ups relative to the full microcircuit model (M0: 1.57 ± 0.02; M50: 1.99 ± 0.03; M100: 2.34 ± 0.03; reduced: 4.88 ± 0.05, *p* < 0.05; *p* < 0.05; **Fig. 3D**).

We next assessed whether these methodologies could be equally applied to a model of a different type of microcircuit, using our layer 5 human microcircuit models (**Fig. 4A**), and found comparable speed-ups (M0: 1.38 ± 0.01; M50: 1.64 ± 0.01; M100: 1.88 ± 0.01; Reduced: 5.21 ± 0.05; *p* < 0.05; **Fig. 4B**). The M50 and reduced models generated unchanged baseline and response spiking (**Fig. 4C**) compared to the M0 models (baseline, M0: 3.24 ± 0.06 Hz, M50: 3.27 ± 0.05 Hz, Reduced: 3.19 ± 0.06 Hz; response, M0: 10.85 ± 0.61 Hz, M50: 11.10 ± 0.64 Hz, Reduced: 10.80 ± 0.73 Hz; **Fig. 4D**). By comparison the M100 models differed in spike rates as before (baseline: 3.41 ± 0.06 Hz, *p* < 0.05, Cohen’s *d* = 2.6; response: 12.29 ± 0.89 Hz, *p* < 0.05, Cohen’s *d* = 1.8; **Fig. 4D**).

**Figure 4.**
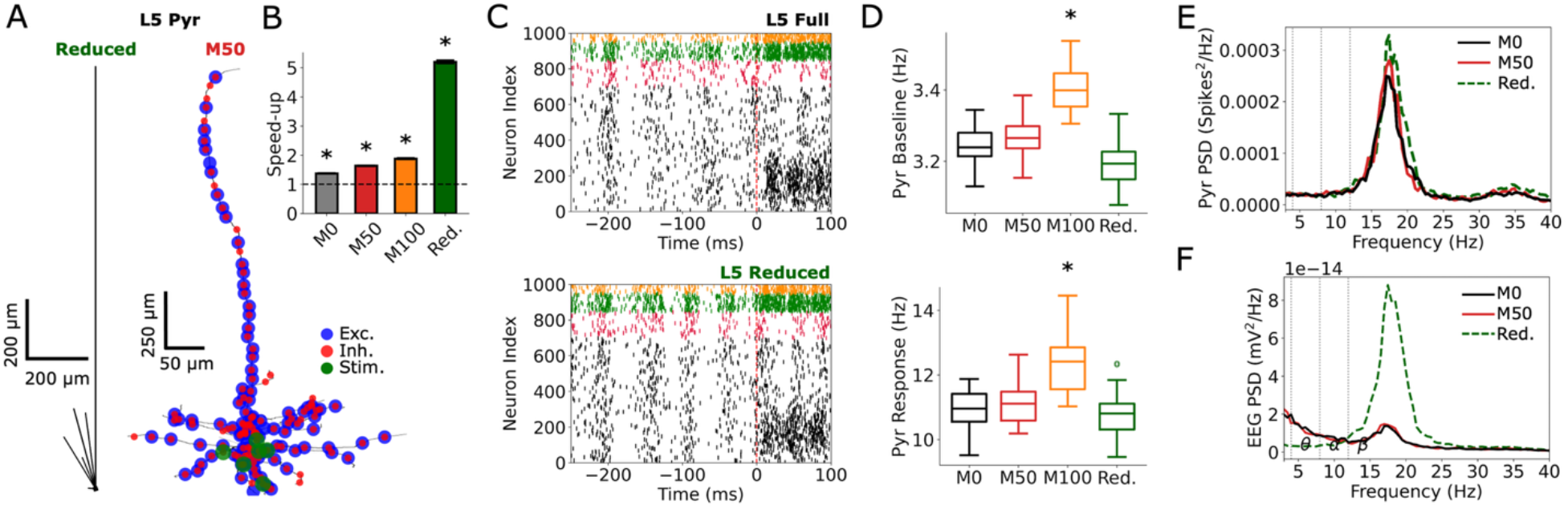
Human L5 microcircuits show similar reduction limits to L2/3 microcircuits. **A**. Human L5 Pyr neuron morphological reduction (left) and example synapse locations resulting from 50 μm synapse merging (right). **B**. Mean speed-ups relative to the full L5 microcircuit model (4.5s-long simulation, without EEG recordings), for the full microcircuit with synapses merged across different spatial resolutions, and the reduced morphology microcircuit (n = 20 microcircuits each). **C**. Example raster plots for full (top) and reduced (bottom) L5 microcircuits. **D**. Baseline (top) and response (bottom) spike rates of Pyr neurons for merged-synapses and reduced microcircuits. **E**. Mean spikes PSD of Pyr neurons for merged-synapses and reduced microcircuits. **F**. Mean PSD of simulated EEG for merged-synapses and reduced microcircuits. Asterisks denote significant differences (*p* < 0.05; Cohen’s *d* > 1) compared to the full microcircuit.

Similarly to the L2/3 cortical microcircuit models, the reduced L5 microcircuit model had differences in spectral properties of spiking and EEG, whereas the M50 models matched well with the M0 model (no differences found). The reduced model exhibited enhanced β (12-30 Hz) power in spike PSD (M0: 0.0012 ± 0.0001 spikes^2^; Reduced: 0.0015 ± 0.0002 spikes^2^, *p* < 0.05, Cohen’s *d* = 1.6; **Fig. 4E**), and in EEG PSD (0.0731 ± 0.0103 nV^2^ vs. 0.3721 ± 0.0547 nV^2^, *p* < 0.05, Cohen’s *d* = 7.4; **Fig. 4F**) that could be quantified by changes to both the aperiodic component (offset: -13.11 ± 0.25 a.u. vs. -15.54 ± 0.16 a.u., *p* < 0.05, Cohen’s *d* = -6.5; exponent: 1.19 ± 0.20 vs. -0.03 ± 0.13, *p* < 0.05, Cohen’s *d* = -7.1) and the periodic component (β peak: 0.60 ± 0.18 a.u. vs. 1.15 ± 0.35 a.u., *p* < 0.05, Cohen’s *d* = 1.9).

Lastly, we assessed the preservation of altered spiking and EEG power spectra in reducing a disease cortical microcircuit model, using our previous major depressive disorder (MDD) layer 2/3 microcircuit models, when reducing SST interneuron inhibition (**Fig. 5A**). The M50 and reduced MDD models preserved baseline and response Pyr neuron spike rates compared to the M0 MDD model (baseline, M0: 1.22 ± 0.04 Hz, M50: 1.23 ± 0.06 Hz, Reduced: 1.18 ± 0.04 Hz; response, M0: 2.41 ± 0.50 Hz, M50: 2.57 ± 0.64 Hz, Reduced: 2.36 ± 0.35 Hz, **Fig. 5B**).

**Figure 5.**
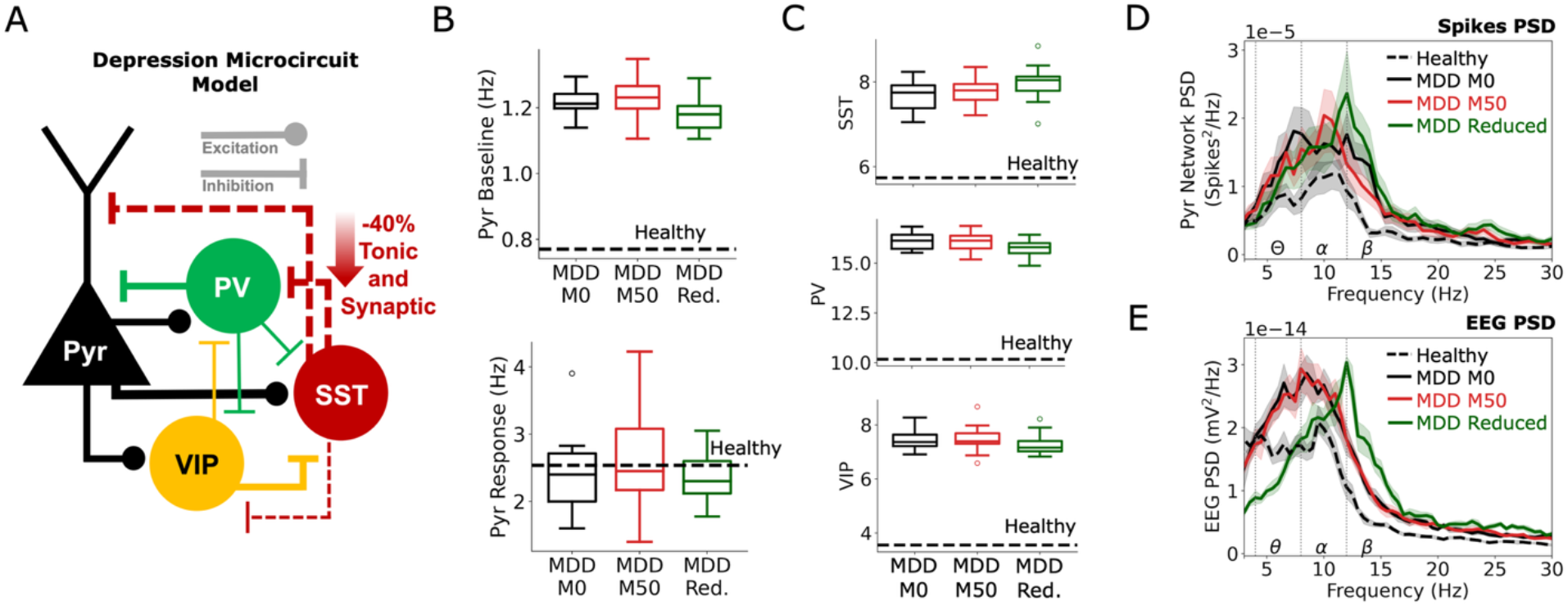
Preservation of activity and EEG effects of depression mechanisms for merged-synapses vs reduced L2/3 microcircuits. **A.** Schematic of the depression (MDD) microcircuit model (Yao et al 2022). **B.** Baseline (top) and response (bottom) spike rates of Pyr neurons for synapse-merged and reduced depression microcircuits (n = 20 microcircuits each). The dashed lines indicate the mean values for healthy full microcircuits. **C.** Baseline spike rates of the different interneuron types for merged-synapses and reduced MDD microcircuits. **D.** Mean spikes PSD of Pyr neurons for merged-synapses and reduced MDD microcircuits. **E.** Mean PSD of simulated EEG for merged-synapses and reduced MDD microcircuits (n = 10 microcircuits each).

Interneuron pre-stimulus spike rates were similarly preserved in the MDD models (SST, M0: 7.67 ± 0.35 Hz, M50: 7.77 ± 0.32 Hz, Reduced: 7.97 ± 0.36 Hz; PV, M0: 16.11 ± 0.41 Hz, M50: 16.08 ± 0.47 Hz, Reduced: 15.76 ± 0.39 Hz; VIP, M0: 7.45 ± 0.34 Hz, M50: 7.46 ± 0.44 Hz, Reduced: 7.26 ± 0.35 Hz, **Fig. 5C**).

The spiking power spectra of the MDD M50 models matched well with the MDD M0 model, and we did not find any differences in θ, α, or β power (**Fig. 5D**). However, the reduced models did show decreased θ power (5.30 × 10^5^ ± 1.50 × 10^5^ spikes^2^ vs. 4.02 × 10^5^ ± 1.47 × 10^5^ spikes^2^, *p* < 0.05, Cohen’s *d* = -0.84) and increased β power (8.63 × 10^5^ ± 1.84 × 10^5^ spikes^2^ vs. 1.08 × 10^4^ ± 3.12 × 10^5^ spikes^2^, *p* < 0.05, Cohen’s *d* = 0.84). For the EEG power spectra, the M50 model matched well with the M0 model (no differences found). However, the reduced model exhibited decreased θ and increased β spectral powers (θ: 0.0940 ± 0.0128 nV^2^ vs. 0.0496 ± 0.0054 nV^2^, *p* < 0.05, Cohen’s *d* = -4.3; β: 0.0689 ± 0.0050 nV^2^ vs. 0.0993 ± 0.0098 nV^2^, *p* < 0.05, Cohen’s *d* = 3.7; **Fig. 5E**), which could be further characterized by changes in the aperiodic component (offset: -13.24 ± 0.08 a.u. vs. -13.82 ± 0.09 a.u., *p* < 0.05, Cohen’s *d* = -6.4; exponent: 0.90 ± 0.06 vs. 0.40 ± 0.07, *p* < 0.05, Cohen’s *d* = -7.4).

## Discussion

In this work, we tested recent analytical morphological reduction and synapse merging techniques on detailed models of human cortical microcircuits. We found that, despite preserving input and output properties of individual neurons, morphological reduction altered power spectral properties of microcircuit spiking and EEG dynamics, whereas synapse merging alone preserved these features. Thus, synapse merging on the full morphologies alone could provide a computational speed up without sacrificing model accuracy. These reduced models can serve in speeding up microcircuit simulations to overcome challenges in optimizations of network properties or more complex cortical functions.

The impact of morphological reduction on spiking dynamics was less pronounced than on EEG dynamics, but in both cases involved increased spectral power of the primary peak. Given that input and output properties of individual neurons were preserved with the morphological reduction method^8^, this effect emerged from the altered spatial spread of synapses. The effect was supported by similar changes seen when we merged synapses in the full morphological models at intervals of 100 μm or more. Because the merging method does not account for dendritic non-linearities^8^, merged synapses and excitatory drives (i.e. Ornstein-Uhlenbeck noise) during coordinated recurrent activity alters local dendritic potentials resulting in a different engagement of dendritic non-linear mechanisms such as NMDA through supra-linear summation of inputs, and *IH* through post-inhibitory rebound dynamics, leading to spiking inaccuracies and enhanced peak power. Accordingly, we did have to partially compensate for this in the morphologically reduced models, by reducing excitatory drive in some cell types, particularly for PV interneuron models, which had higher baseline drives, and applying a surface area compensation factor to *GH* and *Gtonic*.

The more pronounced effect on EEG in the morphologically reduced models was due to the sensitivity of dipole moments to a loss of dendritic branching and elongated apical dendrites, which directly alter the spatiotemporal transmembrane source-sink currents from which the dipole moments are computed^13^. One possible alternative approach to better preserving the EEG may be through a hybrid approach, by computing the EEG post hoc from the simulated morphologically reduced microcircuit activity using a passive template Pyr neuron morphology^22^. As well, toolboxes such as DendroTweaks^23^, which incorporates analytical reduction tools including the ones in this work^8,14^, may be useful for manually fine-tuning the level of apical branching in the reduced pyramidal neuron morphologies to better preserve the EEG.

Despite the shortcomings of the morphologically reduced microcircuit models, both this reduction method and the synapse merging method are useful to facilitate further investigations. The methods can facilitate parameter search for cortical microcircuits, and assessing parameter degeneracy^24^, sensitivity^25^ and identifiability^26^. The methods can also speed up machine-learning algorithms^17,27^ for biophysical microcircuit models, which optimize the connectivity parameters of the model to better capture target outputs or perform specific tasks. In particular, the faster simulations enabled by morphological model reductions may serve in cases of more demanding optimizations, as a first step of the fitting, and be converted back to the full morphological model to further refine the fitted circuitry parameters. Faster simulations can also serve in computational studies that require longer simulated data^28^.

In this work, we tested performance of existing detailed microcircuit models, and their non-linear dendritic mechanisms (*INMDA, IH*) were key contributors to challenges in preserving spiking and EEG dynamics. Future studies should assess other dendritic non-linearities, such as dendritic sodium and potassium channels to promote spike backpropagation^29^ or calcium channels to promote dendritic spiking^30,31^ to refine our findings. In addition, future studies can refine our findings by testing the reduction methods on multi-layered microcircuits that include both L2/3 and L5, to test preservation of dynamics and signal processing across layers^5,17^, as well as test the preservation of cortical response to more complex stimuli and tasks^32^. Future studies may further increase the simulations speed up by using the reduced models methods together with efficient compute simulators such as CoreNEURON^33^.

## Methods

### Detailed models of human L2/3 and L5 cortical microcircuits

We used previous detailed models of human L2/3^1^ and L5^21^ cortical microcircuits. Both models included 1000 neurons of four key cell types (L2/3: 80% Pyr, 5% SST, 7% PV, and 8% VIP; L5: 70% Pyr, 15% SST, 10% PV, and 5% VIP; consistent with proportions in human cortical data^34,35^), with positioning corresponding to their respective volumes (L2/3: 500x500x950 µm^3^; L5: 500x500x700 µm^3^). The neuron models were constrained with data of human neuronal firing^36,37^ and synaptic properties^38–42^ where available, and used human neuron morphologies^37^ with conductance-based ion channels^31^ and probabilistic synapses with short-term dynamics^30^. Excitatory synapses were AMPA and NMDA, and inhibitory synapses were GABAA. Simulations were run using NEURON 7.7^43^ and LFPy 2.0.2 (Python 3.7.6)^13^ on SciNet parallel computing^44^.

### Microcircuit baseline spiking activity

For both L2/3 and L5 models, we drove baseline activity with background excitatory inputs of Ornstein Uhlenbeck (OU) point processes placed throughout the dendritic arbor of each cell^1,21,45^. Specifically, these were placed midway along the length of each dendritic arbor, and for L2/3 and L5 Pyr neuron models, we placed 5 additional OU processes at 10%, 30%, 50%, 70%, 90% of the apical length. We scaled up the mean and standard deviation of each OU conductance exponentially with relative distance from soma (ranging from 0 to 1) to normalize their effect.

For microcircuit baseline spiking we also computed the power spectral density (PSD) of Pyr neuron population spiking by first converting the spike times into binary spike train vectors and then summing the binary spike train vectors across all Pyr neurons. The PSD was then computed from the summed spike train vectors using Welch’s method^46,47^ from the SciPy python module with 1.5 s windows. We quantified the resulting power area-under-the-curve in the θ (4 - 8 Hz), α (8 – 12 Hz), and β (12 – 30 Hz) frequency bands.

### Morphological reductions

To reduce the neuron model morphologies we used the *Neuron_Reduce* toolbox (*subtree_reductor* function), which preserves dendritic transfer impedance to the soma^8^. First, the model reductions parameters were minimally tuned to optimize replication of current-step injection voltage and spiking features. In all cases, we used automatic segmentation (total_segments_manual = -1) and chose a steady-state electrical tuning frequency (reduction_frequency = 0), except the L5 Pyr neuron model, for which we set the reduced model to have a total of 30 segments, a larger number of segments than the default method generated, to improve the reduction fit. Additionally, to improve the reduction fits we found it necessary to implement a conductance and membrane capacitance scaling factor (*f*), described in previous studies^9,10^. This factor scaled the dendritic membrane capacitance and ion channel conductance values in each reduced segment based on the ratio of change in the surface area from complex to reduced segments. The total numbers of sections and segments for the complex and reduced versions of each cell type can be found in **Table 1**. Once reduced, we performed spatial transformations to preserve the initial layout of the morphologies, to improve preservation of EEG. This included rotating the reduced morphologies such that the longest dendritic section of each neuron was vertically oriented and then translating the morphologies to ensure each soma was positioned in its original location. We additionally applied a z-axis translation correction to prevent the straightened dendrites in the reduced models from crossing the pia mater.

**Table 1.** Number of single-cell sections and compartments and simulation times before and after reduction.

| Cell type | Number of full compartments | Number of reduced compartments | $G_{OU}$ Factor |
| --- | --- | --- | --- |
| L2/3 Pyr | 382 | 63 | 0.94 |
| L2/3 & L5 SST | 231 | 44 | L2/3: 0.96<br>L5: 1.00 |
| L2/3 PV | 274 | 57 | 0.65 |
| L2/3 & L5 VIP | 178 | 14 | L2/3: 0.96<br>L5: 1.00 |
| L5 Pyr | 351 | 32 | 0.91 |
| L5 PV | 157 | 65 | 0.52 |

### Synapse merging

To re-position and merge synapses, we again used the methods available through the *subtree_reductor* function. These methods first move synapses from the complex morphological model segments to the equivalent reduced segments and then merge all synapses of the same type on each reduced segment to further reduce computational load. To merge synapses at spatial intervals without morphological reduction, we customized these methods to re-position and merge synapses onto the full morphologies following specified spatial intervals. For example, if merging synapses onto every 50 μm location from the soma, we would first identify each 50 μm dendritic interval from the soma and then merge all synapses to their corresponding nearest identified locations.

### Ornstein-Uhlenbeck noise re-positioning and tuning

We applied the same mapping of complex to reduced segments to re-position and re-initiate the OU point processes onto their equivalent reduced segments. Following reductions, we found that further tuning of OU conductance values (*GOU*) was needed to preserve microcircuit spiking, particularly for PV interneurons (see *GOU* factors in **Table 1**).

### Microcircuit response spiking activity

We used the stimulus paradigm developed previously for the L2/3 model^1^, which used excitatory AMPA/NMDA synapses with the same synaptic dynamics as the cortical excitatory synapses. We stimulated the basal dendrites of 55 Pyr neurons, with 2–4 ms delay post-stimulus and a conductance of 4 nS. We also stimulated 35 PV interneurons with a delay of 2–2.5 ms and a conductance of 2 nS. VIP interneurons were stimulated in two groups and phases: early (65 VIP interneurons, delay = 2–2.5 ms, conductance = 2.8 nS) and late (80 VIP interneurons, delay = 7–12 ms, conductance = 2.2 nS).

For L5 microcircuits, we used the stimulus paradigm developed previously^21^, which applied stimulus orientation selective stimuli to 50% of the Pyr neurons in the circuit, where Pyr neuron with response selectivity near their peak center were stimulated with 45 excitatory synapses, and 30 excitatory synapses for PV and VIP interneurons. Presynaptic inputs were set to spike at 100 Hz over the course of a 100 ms time window.

### Simulated microcircuit EEG and power spectral analysis

We used LFPy to simulate resting-state EEG time series from our microcircuit models^2^, using a four-sphere volume conductor model (representing grey matter, cerebrospinal fluid, skull, and scalp with radii of 79 mm, 80 mm, 85 mm, and 90 mm, respectively)^2^ that assumes homogeneous, isotropic, and linear conductivity. For each sphere, the conductivity was 0.047 S m^−1^, 1.71 S m^−1^, 0.02 S m^−1^, and 0.41 S m^−1^, respectively^2,48^. We computed power spectral density (PSD) using Welch’s method^47^ with 2 s windows, and quantified the resulting power area-under-the-curve in the θ, α, and β frequency bands. We decomposed the EEG power spectra (in the 3 – 30 Hz range) into periodic and aperiodic components using the FOOOF toolbox^49^. The aperiodic component was a 1/*f* function parameterized by vertical offset and exponent parameters. We fitted the periodic oscillatory component with up to 3 Gaussian peaks defined by center frequency, bandwidth (min: 2 Hz, max: 6 Hz), and power magnitude (relative peak threshold: 2, minimum peak height: 0)^2,3^.

### Depression microcircuit models

We used our previous depression microcircuit models^1,7^, with reduced tonic and synaptic conductance of SST interneuron inhibition onto all neuron types in the microcircuit by 40%, in line with postmortem expression studies in depression patients^50^. When reducing tonic inhibition, we scaled the decreased tonic inhibition conductance by the relative contribution of SST interneuron inhibition to total inhibition onto each neuron^1^.

### Statistical tests

We performed Welch’s t-tests to identify differences between complex and reduced models and computed Cohen’s *d* for effect sizes.

## Code availability

All code will be available at https://doi.org/10.5281/zenodo.22800117 as of the date of publication.

## Acknowledgements

EH thanks the Krembil Foundation for funding support.

